# Neurophysiological correlates of processing Agreement and Tense in Arabic

**DOI:** 10.64898/2026.04.10.717434

**Authors:** Ali Idrissi, R. Muralikrishnan

## Abstract

Most syntactic approaches converge on the fact that Tense and Agreement are two different functional categories, although there is less agreement on their exact representation and relative hierarchical order. Cross-linguistic agrammatic data seems to support the difference between Tense and Agreement, with patterns of dissociation reported from agrammatism between them, in which Tense is generally more impaired than Agreement. To examine whether there is evidence for such a dissociation of tense and agreement processing in neurotypical individuals, the present study employed Event-Related brain Potentials (ERPs) to study the real-time comprehension of Modern Standard Arabic sentences. Critical stimulus sentences were of the form Temporal Adverb–Subject–Verb–PP, in which the intransitive verb was in either the past or future tense, and was preceded by a singular or plural subject and an adverb indicating past or future tense. The subject nouns were all human and either masculine or feminine. The verbs either agreed with the subject noun or presented a person, number or gender agreement violation. They also either agreed or showed a mismatch with the temporal frame of the adverb, the latter being a tense violation. Results at the verb showed that both tense and agreement violations yielded a biphasic N400 – P600 effect. We discuss these results in light of previous ERP findings and conclude that despite the putative configurational differences between Tense and Agreement, the processing of the two categories in Arabic may deploy the same underlying cognitive mechanisms.

## 1 Introduction

Crosslinguistically, structural and semantic dependencies among one or more constituents of a sentence tend to be grammatically expressed. For example, the semantic or syntactic properties of a noun, say gender and number, may be overtly expressed on the verbs, determiners or adjectives within the same sentence (Corbett, 2006; Wunderlich, 2015). This phenomenon is commonly known as grammatical agreement. Likewise, in a similar form of agreement, the location of an event or state in time, which may or may not be correlated with a temporal expression in a given sentence, may be overtly expressed on the verb in the form of a Tense morpheme (Comrie, 1985). In either case, there is a systematic covariance between the semantic or syntactic properties of a constituent in a given sentence and the formal and semantic features of another constituent in the same sentence. This is the hallmark of agreement, in its more general sense, whereby agreement, or grammatical feature matching is grammaticalized by means of inflectional morphemes in natural languages.

Theoretical linguistic models have offered analyses of the functional categories of Tense and Agreement to account for their grammatical and structural properties. A famous account that followed the beginning of Chomsky’s Minimalism (Chomsky, 1995) is the Split Inflection Hypothesis (Pollock, 1989). Under this hypothesis, the then traditional Inflection Phrase (IP) node in the syntactic tree is split into separate functional categories with a fixed hierarchical order: highest Complementizer Phrase (CP), then Tense Phrase (TP), followed by Agreement Phrase (AgrP) and, lowest among all, Verb Phrase (VP). In a later version of Minimalism, Chomsky (2000) treats tense as a (semantically) interpretable feature of the category T, while Agr is a feature within T. In other words, TP hosts both the interpretable Tense features and the uninterpretable Agreement features. Within this theory, agreement boils down to an operation whereby the uninterpretable features of T are checked (Chomsky, 2000).

Since the mid-1990s, several behavioral neuropsychological studies have tested the validity of these approaches and the psychological reality of the representations they presuppose (Burchert, Swoboda-Moll, & Bleser, 2005; Friedmann, 2001; Friedmann & Grodzinsky, 1997, 2000; Lee, 2003; Nanousi, Masterson, Druks, & Atkinson, 2006; Varlokosta et al., 2006; Wenzlaff & Clahsen, 2004). Then, as the use of EEG (electroencephalography) technology became more accessible and used in the study of language processing, many researchers started to explore the time course of processing of agreement and tense, and the scalp distribution of their neurophysiological correlates. A relatively large body of results bearing on Agreement processing, mostly in European languages, is now available (Molinaro, Barber, & Carreiras, 2011), including two studies on Arabic (Idrissi, Mustafawi, Khwaileh, & Muralikrishnan, 2021, and Muralikrishnan & Idrissi, 2021). The neural correlates of the processing of tense, also mainly in European languages, have also been investigated, although to a lesser extent than grammatical agreement (Baggio, 2004, 2008). In this paper, we examine the neurophysiological correlates of the processing of Tense and Agreement in Modern Standard Arabic (StA) to shed light on the cognitive mechanisms underlying their processing during online language comprehension.

The paper is organized as follows. Section 2 reviews prior behavioural and neurophysiological research on the representation and processing of Tense and Agreement. Section 3 presents the study and its hypotheses. Section 4 describes the methodology, and Section 5 reports the main results. Section 6 discusses these results and concludes the paper.

## 2 Previous studies

### 2.1 Behavioural studies

Many studies reported data from agrammatic aphasia showing patterns of selective impairment between Tense and Agreement, with Tense morphemes being more impaired than Agreement morphemes. The data obtained from these studies was interpreted as strong evidence not only for the status of Tense and Agreement as separate functional categories, but also for their relative hierarchical order in the syntactic tree, along the lines of Pollock’s (1989) Split-Inflection Hypothesis. Additionally, and since the Tense node is assumed to be higher than Agr in the syntactic tree, this data was taken to indicate that functional nodes higher in the tree are more vulnerable and, as such, (i) should be more likely to be impaired in agrammatic speech, and (ii) can be intact only if lower nodes are. This view, first expressed by Hagiwara (1995), was later mainly associated with the Tree-Pruning Hypothesis (TPH) (Friedmann, 2001, 2005; Friedmann & Grodzinsky, 1997). Within the TPH, Tense is more vulnerable than Agreement: it is affected by the deficit, which is metaphorically viewed as a form of ‘pruning’ the tree. If also follows from this view that, in general, the lower the deficit (or ‘pruning’) in the syntactic tree, the more severe the impairment of functional categories.

Figure 1 shows the deficit site in the tree that leads to the observed tense/agreement dissociations in agrammatic patients (Friedmann, 2005, p.1046).

**Figure 1:**
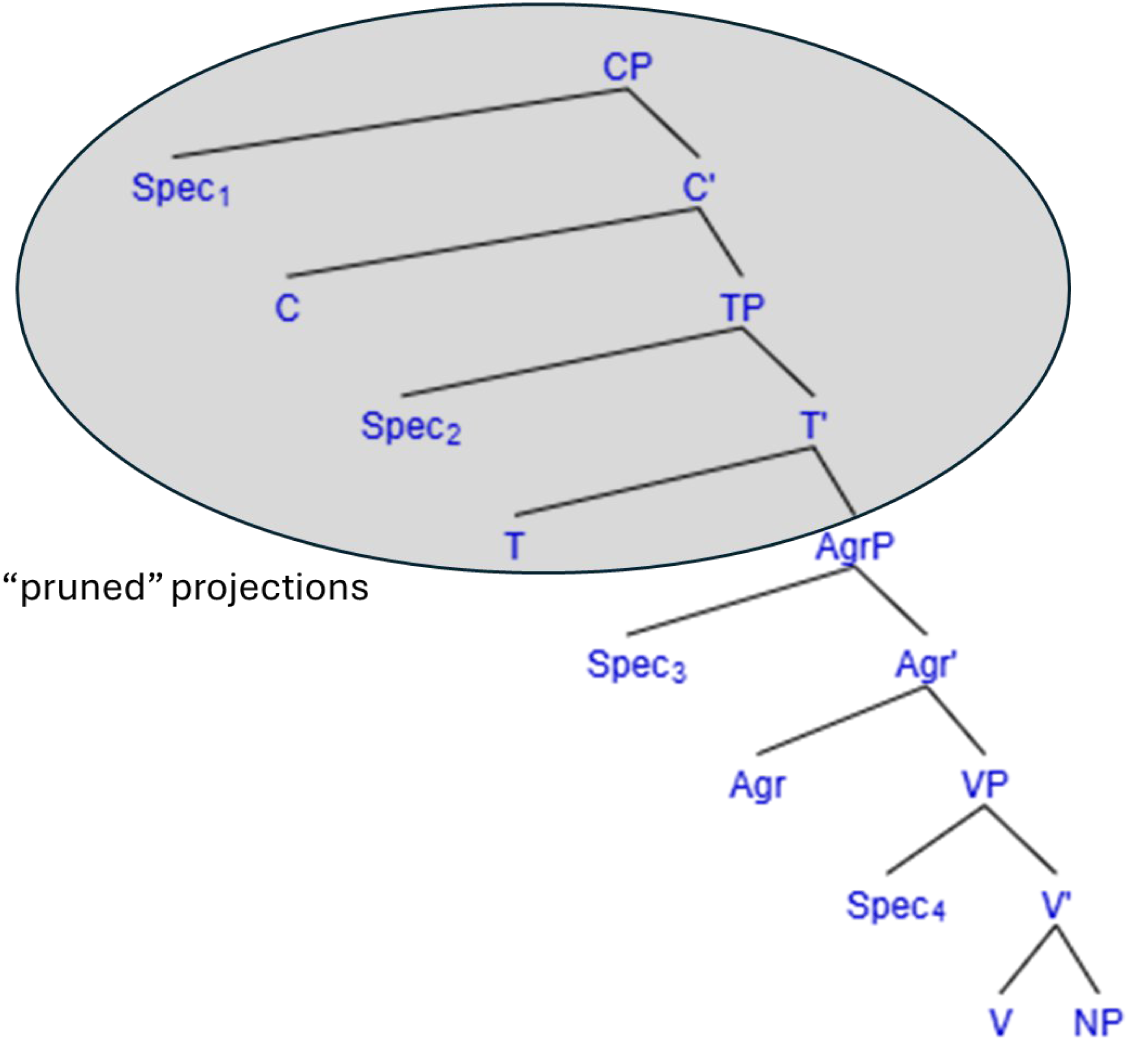
Agrammatic pruned tree showing the pruned projections and site of pruning (Friedmann and Grodzinsky, 1997)

However, various studies question the TPH-based explanation of the dissociation between Tense and Agreement. For example, Lee (2003) reports data from a Korean agrammatic patient that goes against the TPH in that higher functional nodes were more preserved than the lower ones. Other researchers reported the same performance patterns in the production and comprehension of Tense and Agreement, but proposed explanations within more recent versions of the Minimalist theory that do not invoke hierarchical structure (Burchert et al., 2005; Wenzlaff & Clahsen, 2004). Yet, other studies on different languages reported inconsistent data or data that did not show an uncontroversial support for the TPH, essentially because the hierarchical organization of functional categories is not the same across languages (Nanousi, Masterson, Druks, & Atkinson, 2006; Varlokosta et al., 2006).

In a study on German-speaking agrammatic patients, Burchert et al. (2005) provide a good example of inconsistent data. These authors studied 9 German-speaking patients, who they classified in two three subgroups depending on their performance on Tense and Agreement. The first subgroup did not show any dissociation and performed well on both Tense and Agreement. The second group did not show any dissociation either, but they performed poorly on both categories. The third group (3 patients) did show a pattern of dissociation, but in different directions: two performed better on Tense than for Agreement, while one showed the opposite pattern. Burchert et al. (2005) conclude that the purported dissociation between Tense and Agreement is not universal, and when it is observed, it is not unidirectional. The authors propose the Tense Agreement Underspecification Hypothesis (TAUH), which predicts relative impairments of Tense and Agreement in either direction. Importantly, no loss of structure is required in order to account for agrammatic performance on Tense and Agreement, and no automatic impairment of either one or the other functional category is necessary. The TAUH explanation is also inspired by the development in syntactic theory whereby tense and agreement were considered features of different types but features that belonged in the same node.

An explanation along the same lines was given by Fyndanis, Varlokosta, & Tsapkini (2012). These authors obtained data from two Greek-speaking agrammatic individuals. They found that Aspect was significantly more impaired than Tense, and Tense significantly more impaired than Agreement. The authors argue that Aspect and Tense are more vulnerable, not because they are higher in the tree, but rather because they carry interpretable (semantic) features. Agreement, by contrast, is more preserved because it carries an uninterpretable feature, whose processing demand is less than that of Tense (Fyndanis et al., 2012).

Finally, it was noted that the picture of the Tense and Agreement dissociation is complicated by the inherent differences between production and comprehension. In fact, patients were reported to be able to understand the meaning of Tense even when they were unable to produce tensed verbs. In a study on Arabic agrammatism (unpublished), results were obtained from an Arabic-speaking agrammatic patient who showed more impairment on Tense than Agreement in production, but more impairment on Agreement than Tense in comprehension/grammaticality judgement.

The problems raised with the TPH may suggest that any hierarchical representation-based explanation for the fate of Tense and Agreement in agrammatism is likely to fail. The patterns that have been observed, while showing a clear dissociation between the two functional categories, are not unidirectional and do not seem to be completely independent of the task (i.e. input versus output-oriented task).

### 2.2 Neurophysiological studies

The ERP technique has been extensively used since early 1980s in probing the cognitive processes underlying language comprehension because of its excellent temporal resolution, making it particularly well-suited for studying language comprehension in real time. Many previous ERP studies have investigated agreement processing (see Molinaro et al., 2011 for a review), with a handful looking into the processing of tense (Baggio, 2008). Most ERP agreement experiments use the violation paradigm, which consists of grammatical structures and their ungrammatical counterparts. Mean amplitude differences between the different conditions at different scalp sites, and at different latencies from the onset of the critical word are then said to reveal the mechanisms involved in the processing of the feature manipulated.

Three major components of the neural responses have been identified as being associated with the processing of morphosyntactic aspects of language (Friederici, 1995), namely the N400, Left-Anterior Negativity (LAN), and P600. The N400 is a negative-going component, peaking between 300 and 450 ms after the onset of the stimulus, mainly in the centroparietal regions. Associated originally with semantic anomalies (Kutas & Hilyard, 1980: Kutas & Federmeier, 2000; Lau, Phillips, & Poeppel, 2008), its amplitude increases as a function of the increase in the demands of cognitive processing. It has been reported in several ERP studies on agreement processing, and especially in the previous studies on Arabic (e.g., Muralikrishnan & Idrissi, 2021; Idrissi et al., 2021), and is said to also reflect processes related to sentence-level expectancy (Alday, Schlesewsky, & Bornkessel-Schlesewsky, 2017) and prediction errors (Bornkessel-Schlesewsky & Schlesewsky, 2019). LAN (Left Anterior Negativity) is also negative-going and shows the same temporal distribution as the N400 but a different topographical distribution, namely in the left-frontal electrode regions. It is believed to arise in response to morphosyntactic processing over ungrammatical constructions (Osterhout & Mobley, 1995), such as agreement violations. The P600 is a positive-going ERP response, peaking between 500 and 1000 ms and showing more posterior scalp distribution. Amongst others, the P600 has been said to correlate with late syntactic analysis and reanalysis/repair of syntactic anomalies (Bornkessel-Schlesewsky & Schlesewsky, 2008; Osterhout & Mobley, 1995) and more generally with conflict-monitoring processes (van de Meerendonk, Kolk, Chwilla, & Vissers, 2009). Notwithstanding the classical association of these components to different linguistic domains, cross-linguistic research on multiple aspects of language processing in typologically diverse languages has shown that such a one-to-one mapping is untenable.

### 2.3 ERP studies of Agreement

Several ERP studies conducted on agreement in various languages have reported a LAN – P600 pattern in response to subject-verb, determiner-noun and noun-adjective feature mismatches (Barber & Carreiras, 2003, 2005; Mancini, Molinaro, Rizzi, & Carreiras, 2011), but this is by no means the only pattern of effects reported for processing agreement violations. An N400 effect has been reported in several studies on agreement (Muralikrishnan & Idrissi, 2021; Idrissi et al., 2021; Bhattamishra et al., 2021; Zawiszewski, Santesteban, & Laka, 2016; Díaz, Sebastián-Gallés, Erdocia, Mueller, & Laka, 2011) instead of a LAN, and no negativity effect ensued at all in some studies (Gulati et al., 2024; Alemán Bañón and Rothman, 2016; Frenck-Mestre et al., 2009; Nevins et al., 2007). In other words, a quick look at a comprehensive overview of cross-linguistic agreement studies (Gulati, Muralikrishnan & Choudhary, 2024) shows that the processing of agreement features elicit a diversity of effects, and that no single pattern of effects can be said to index agreement violations across languages and structures. Rather, Muralikrishnan & Idrissi (2021) have posited that the diversity of cross-linguistic ERP effects for agreement processing can be accounted for by considering the relative salience of each feature in a given language.

### 2.4 ERP studies of Temporality

Most of the early studies that touch upon the processing of tense dealt with the ERP correlates of the processing of regular vs. irregular verb forms, mostly in English. However, a few later studies investigated the ERP responses to mismatches between tense morphology on the verb and the temporal frame expressed by the temporal adverbs or temporal adjunct clauses. Note that the notion of tense violation is not as straightforward as agreement violation, since it is a semantic notion (see Baggio 2004, 2008). In one of the first studies on tense violations, Kutas & Hillyard (1983) reported an N400 like effect in English. Osterhout and Nicol (1999) reported a P600 effect for tense violations involving modal constructions in English (can fly/*flying). Fonteneau et al. (1998) reported a negativity followed by a late positivity in French tense violations. Similarly, Allen, Badecker & Osterhout (2003) reported a biphasic N400 – P600 for tense violations in English, whereas Steinhauer & Ullman (2003) reported LAN – P600 to tense violations such as ‘Yesterday, I sailed/*I sail’. Baggio (2004, 2008) reported LAN-P600 for tense violations in Dutch.

## 3 Current Study

While the behavioral neuropsychological data often showed a dissociation between Tense and Agreement (although not in the sense argued for by the TPH), which may suggest a processing difference between the two functional categories, the ERP data do not seem to show much difference. Without making any claims about the possible difference between behavioral and ERP evidence, we propose to investigate the ERPs of Tense and Agreement violations using stimuli in StA, a language in which the two categories are expressed differently in the morphology: tense seems to be encoded templatically, and agreement is marked with affixes. We hypothesize that if the underlying cognitive processes involved in the processing of tense and agreement are sensitive to the nature of how each category is morphologically encoded and the morphosyntactic and semantic differences between the two categories, we should observe qualitative ERP differences between the processing of tense and agreement in Arabic. Alternatively, if tense and agreement are processed in the same way during online comprehension, we should observe the same ERP effects for the processing of tense and agreement regardless of how they are morphologically encoded.

## 4 Methods

### 4.1 Participants

Twenty-eight participants, mostly students, residing in the UAE, participated in the experiment after giving informed consent (nine female; mean age: 22.14 years; age range: 19 – 27 years). Five participants had to be excluded from the final data analyses because of EEG artefacts and/or too many errors on the behavioral control task. All participants were right-handed native-speakers of Arabic from different parts of the Arab world, with normal or corrected-to-normal vision and normal hearing.

### 4.2 Materials

The experiment consisted of three critical conditions of the form adverb-subject-verb-prepositional phrase. Table 1 provides an overview of the conditions, and Table 2 illustrates them with actual examples. The critical position was at the verb, which differed between the critical conditions as follows. The stimulus verb was either fully acceptable, or it posed a violation of agreement with the subject. The violation was either in person, number, or gender; never in a combination of two features; or else a violation of tense whereby the morphology of the verb clashes with the temporal window defined by the sentence-initial temporal adverb.

**Table 1:** Conditions.

| Factor | Level |
| --- | --- |
| CT: Condition | ACP: Acceptable |
|  | VAG: Violation of agreement |
|  | VTN: Violation of tense |

**Table 2:**
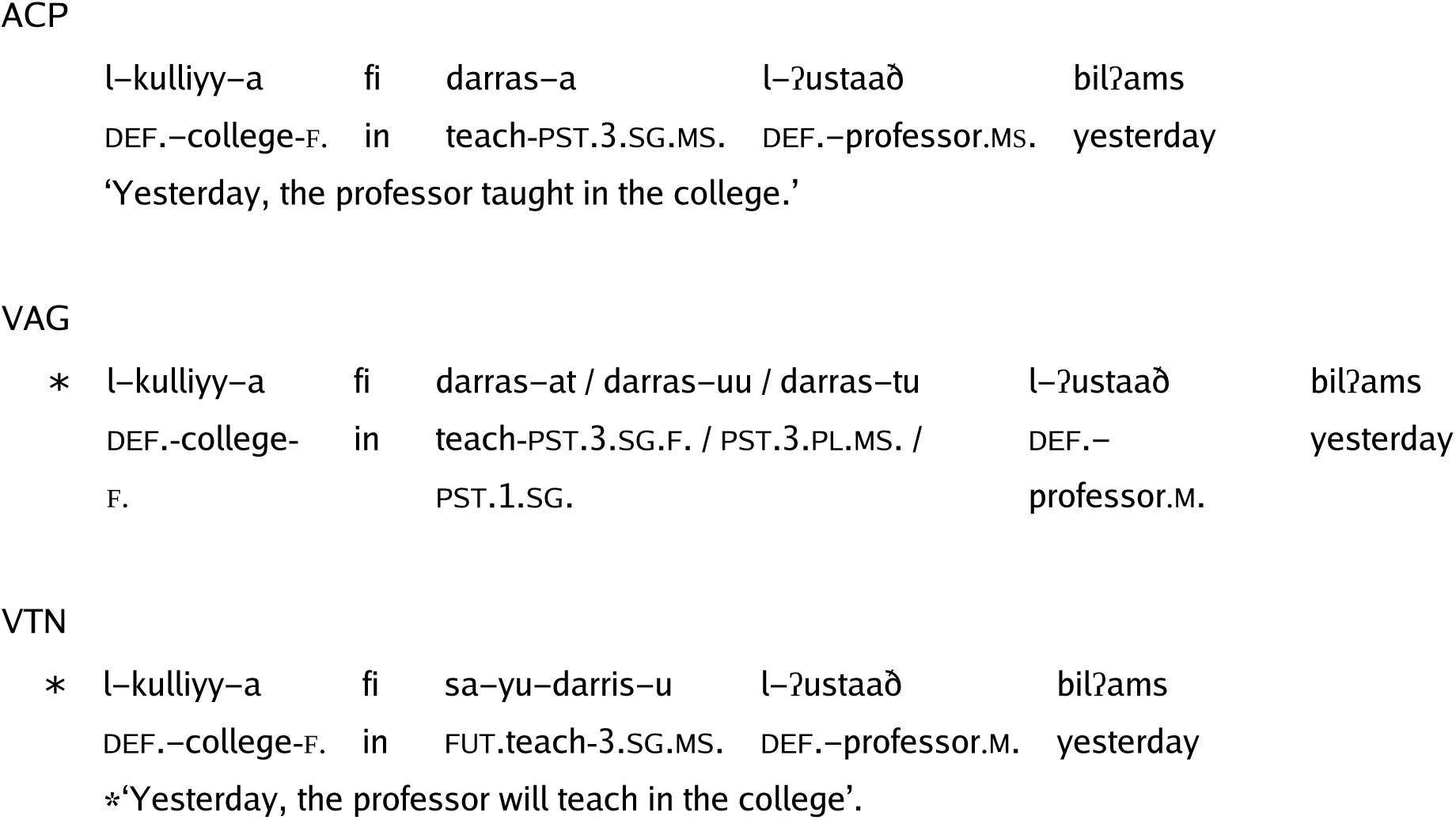
The critical stimulus sentences. The conditions are coded as ACP (acceptable), VAG (violation of agreement) and VTN (violation of tense).

|  |  |  |  |  |
| --- | --- | --- | --- | --- |
| l-kulliyy-a | fi | darras-a | l-ʔustaað | bilʔams |
| DEF.-college-F. | in | teach-PST.3.SG.MS. | DEF.-professor.MS. | yesterday |
| **VAG** |  |  |  |  |
| * l-kulliyy-a | fi | darras-at / darras-uu / darras-tu | l-ʔustaað | bilʔams |
| DEF.-college-F. | in | teach-PST.3.SG.F. / PST.3.PL.MS. / PST.1.SG. | DEF.- | yesterday |
|  |  |  | professor.M. |  |
| **VTN** |  |  |  |  |
| * l-kulliyy-a | fi | sa-yu-darris-u | l-ʔustaað | bilʔams |
| DEF.-college-F. | in | FUT.teach-3.SG.MS. | DEF.-professor.M. | yesterday |

\*‘Yesterday, the professor will teach in the college’.

Subject nouns were always human. As a first step, 120 intransitive verbs were used to build acceptable sentences with singular subject nouns, half of which were masculine and the other half feminine. The verbs were in the past tense and the adverb was always *yesterday*. These sentences were then adapted to generate a further 120 acceptable sentences such that the adverb was now *tomorrow*, the subject nouns were plural, and the verb showed future tense. Then, three types of agreement violation conditions were generated from the acceptable sentences. Again, here the violations involved person, number or gender separately (that is, never in combination). Finally, two tense violation conditions were created such that the verb bore the past tense morphology with the adverb *tomorrow* or future tense morphology in the context of the temporal adverb *yesterday*.

The critical sentences were distributed into two unique sets according to a Latin square, such that each list contained 120 acceptable sentences, 120 tense violation sentences, and 120 agreement violation sentences (40 person-agreement violations, 40 number agreement violations, and 40 gender agreement violations). Half of each type of sentence in each set contained a singular noun and was in the future tense; the other half contained a plural noun, and was in the past tense. The resulting 360 sentences were then interspersed with fillers of various types, including semantic violations and relative clause constructions. The result is that, overall, each stimulus list ended up with an equal number of acceptable and violation sentences, an equal number of sentences with a masculine or feminine, singular or plural noun, and an equal number of sentences in the past and future tenses, which led in the end to a total of 528 sentences per stimulus list. The two lists were each pseudo-randomized thrice to obtain six stimulus lists, one of which was used for every participant. The presentation of the randomized lists was counterbalanced across participants. The following abbreviations are used in the morphological glossing of the example stimuli in Table 2: PST. = past, FUT. = future, 1 = first person, 3 = 3^rd^ person, SG. = singular, PL. = plural, MS. = masculine, F. = feminine, DEF. = definite. The Arabic text is read from right to left.

### 4.3 Procedure

The experiment was performed in the EEG laboratory of the Department of Linguistics at the United Arab Emirates University in Al Ain. The methods and procedure employed in the experiment were in accordance with the University Ethics Committee regulations and followed the guidelines of the Helsinki declaration. Upon their arrival at the lab, the participants filled in an Edinburgh-Handedness questionnaire in Arabic. Only dominant right-handers were accepted. Participants were given printed information about the experiment and instructions relative to the task they had to perform. Stimuli were presented using the Presentation software (www.neurobs.com) that recorded, among other things, the trial number, reaction time and the key or button used to register answers. The brightness and contrast settings of the monitor were maintained the same for all the participants.

After setting up the electrode cap, the participant moved to a soundproof chamber, where they were seated on a comfortable chair and were requested to avoid abrupt and drastic movements, especially of the head. Then the so-called ‘resting EEG’ was recorded for possible subsequent frequency-based EEG analyses, where the participant had to sit still for two minutes with no specific task to perform. Two more minutes of resting EEG was recorded, but now the participants had to close their eyes. After a short pause, the experimental session commenced, which consisted of a short practice session followed by the actual experiment.

The structure of an experimental trial was as follows. The flat-screen LCD monitor was clear before the trial commenced. A fixation asterisk was shown in the center of the screen for 500 ms, after which the screen became blank for 100 ms. Then, the rapid serial visual presentation of the stimulus sentence started. Each word appeared in the center of the screen and remained for 600 ms, after which the screen became blank for 100 ms before the appearance of the next word. When they consist of two separate words, the preposition and its object noun phrase, the prepositional phrases were presented for 750 ms. After the last word of the stimulus sentence was presented, the screen was blank for 500 ms. Following this, a pair of smileys appeared on screen, which prompted the participant to judge the acceptability of the sentence that just preceded. After a maximum of 2000 ms or after a button press, whichever comes earlier, the screen became blank again, for 500 ms. A time-out was registered when no button was pressed within 2000 ms. Then, a probe word appeared in the middle of the screen for a maximum of 2000 ms, within which the participant had to detect whether the word was present in the preceding stimulus sentence or not. When no button was pressed within 2000 ms, a time-out was registered. At the end of the trial, the screen became blank for 1500 ms (inter-stimulus interval) before the next trial started.

Before the actual experiment began, there was a short practice consisting of twelve trials, which helped participants to get used to the task and feel comfortable with the pace of the trials and the blinking regime. For a given participant, none of the experimental stimuli occurred in their practice phase. The task was identical to that of the experiment phase. The EEG of the participants was not recorded in this phase.

In the main phase of the experiment, one of the six sets of materials as mentioned earlier was chosen to be presented in 12 blocks of 44 trials each. Given the use of a violation paradigm, an acceptability judgement task followed the presentation of each stimulus sentence, which required a ‘yes’ or ‘no’ answer. In addition, in order to ensure that participants would process the sentences attentively, a probe word detection task followed the acceptability judgement task. The probe task was constructed in such a way that an equal number of trials required ‘Yes’ or ‘No’ as answers. If the probe word was one of the words that occurred in the preceding stimulus sentence, this required a ‘yes’, whereas if it was novel, it required a ‘no’. Crucially, the word position from which the probe word was chosen was equiprobable across the experiment as well as within each condition, which meant that participants had to be very attentive throughout stimulus presentation for them to perform the task correctly. There was an equal number of probe words that required ‘Yes’ or ‘No’ as answers in each block. In order to counterbalance for any right-dominance effects, we had the ‘Yes’ button on the right side for half the number of participants and on the left side for the other half. The ‘Yes’ button being on the right or left was also counterbalanced across the stimuli sets. There was a short pause between blocks. Resting EEG was again recorded at the end of the experimental session.

### 4.4 EEG recording, pre-processing and statistical analysis

The EEG was recorded by means of 25 AgCl-electrodes fixed at the scalp by means of an elastic cap (Easycap GmbH, Herrsching, Germany). AFZ served as the ground electrode. Recordings were referenced to the left mastoid but re-referenced to the average of linked mastoids offline. The electrooculogram (EOG) was monitored by means of electrodes placed at the outer canthus of each eye for the horizontal EOG, and above and below the participant’s right eye for the vertical EOG. Electrode impedances were kept below appropriate levels such as to ensure a good quality signal with minimal noise. All EEG and EOG channels were amplified using a Brain Amp amplifier (Brain Products GmbH, Gilching, Germany) and recorded with a digitization rate of 250 Hz. The EEG data thus collected was pre-processed for further analysis using a band pass filter that passed signals in the frequency range 0.3 Hz to 20 Hz. These filter settings are sufficiently broad to include language-related ERP activity that is typically in the frequency range of about 0.5–5 Hz (Delorme, 2023; Roehm et al., 2002) and have been employed in several previous cross-linguistic ERP studies on language processing in diverse languages. For the purposes of visualization, an 8.5 Hz low-pass filter was further applied on the data for achieving smoother ERP plots.

ERPs were calculated for each participant from 200 ms before the onset of the verb until 1200 ms after onset (so -200 ms to 1200 ms). These were averaged across items per condition per participant before computing the grand-average ERPs across participants per condition. Only those trials in which the participants performed the probe detection task correctly were considered for the analysis. The statistical analysis of the ERP data was performed by fitting linear mixed-effects models in R (Version 4.5.2, R Core Team, 2025) using the lme4 package (Version 2.0.0, Bates et al., 2015). Interactions were resolved by computing estimated marginal means on the response scale using the emmeans package (Lenth & Piaskowski, 2025) with the mvt multivariate adjustment applied for multiple comparisons. The statistical models included the fixed factor Condition (with three levels: Acceptable, Violation of Agreement, Violation of Tense), as well as the topographical factor regions of interest (ROI). Separate models were fitted for lateral ROIs and midline ROIs for mean amplitude values per time-window per condition. The 4 lateral ROIs were defined as follows: LA, comprised of the left-anterior electrodes F7, F3, FC5 and FC1; LP, comprised of the left-posterior electrodes P7, P3, CP5 and CP1; RA, comprised of the right-anterior electrodes F8, F4, FC6 and FC2; and RP, comprised of the right posterior electrodes P8, P4, CP6 and CP2. Each of the 6 mid-line electrodes FZ, FCZ, CZ, CPZ, PZ and POZ constituted an individual midline ROI of the same name respectively. Furthermore, rather than performing a traditional subtraction-based baseline correction, the mean 200 ms pre-stimulus baseline amplitude (−200 to 0 ms) in each data epoch was included as a covariate (after scaling) in the statistical model. This was done in order to account for and regress out potential contributions of differences in the baseline period in the statistical analysis (Alday, 2019). However, we do not interpret effects involving pre-stimulus amplitudes, because these did not form part of our hypotheses. Further, this is in line with the fact that “we can include additional covariates as controls without further interpreting those covariates” (Alday, 2019, p.9). The contrasts for the categorical factors used sum contrasts, such that coefficients reflect differences to the grand mean (Schad et al., 2020). Except where specified otherwise, the random effects structure was maximal (Barr et al., 2013), with random intercepts for participants, and by-participant random slopes for the effect of Condition. Following modern statistical recommendations, we do not use the term ‘statistically significant’ or its variants based on p-value thresholds (Wasserstein et al., 2019), but report precise p-values as continuous quantities (e.g., p = 0.06 rather than p < 0.08), unless a value is “below the limit of numerical accuracy of the data”, in which case, we report it as p < 0.001 (Amrhein et al., 2019, p.206). Further, we supplement the p-values by transforming them into s-values (Shannon information, surprisal, or binary logworth) and report s = – log2(p), which provides a nonprobability measure of the information provided by a p-value on an absolute scale (Shannon, 1948; Greenland, 2019). In other words, “the s-value provides a gauge of the information supplied by a statistical test” and has the advantage of providing “a direct quantification of information without” requiring prior distributions as input (Rafi & Greenland, 2020, p.6). The analysis code and full model outputs are available as R notebooks in the data analysis repository online.

## 5 Results

### 5.1 Behavioral data

The mean acceptability ratings for the stimuli, as well as the probe detection accuracy for the critical conditions, shown in Table 3, were calculated using the behavioral data collected during the experiment. Only those trials in which the acceptability judgement task following each trial was performed (i.e., not timed out) were considered for the analysis. Further, the acceptability data presented here pertain only to those trials in which the participants performed the probe detection task correctly. Acceptability was highest for the acceptable condition, whereas it was the lowest for the tense violation condition. The overall accuracy was very high in all conditions.

**Table 3:** Mean acceptability ratings and probe detection accuracy.

| Condition | Acceptability % | SD | Accuracy % | SD |
| --- | --- | --- | --- | --- |
| ACP | 88.6 | 31.8 | 93.9 | 24.0 |
| VAG | 30.9 | 46.2 | 92.7 | 26.0 |
| VTN | 20.1 | 40.1 | 91.2 | 28.4 |

Figure 2 shows raincloud plots (Allen et al., 2021) of the behavioral acceptability judgements. Panel A shows the by-participant variability of acceptability ratings, with the individual data points representing the mean by-participant acceptability of each condition. Panel B shows the by-item variability of acceptability ratings, with the individual data points representing the mean by-item acceptability of each condition.

**Figure 2:**
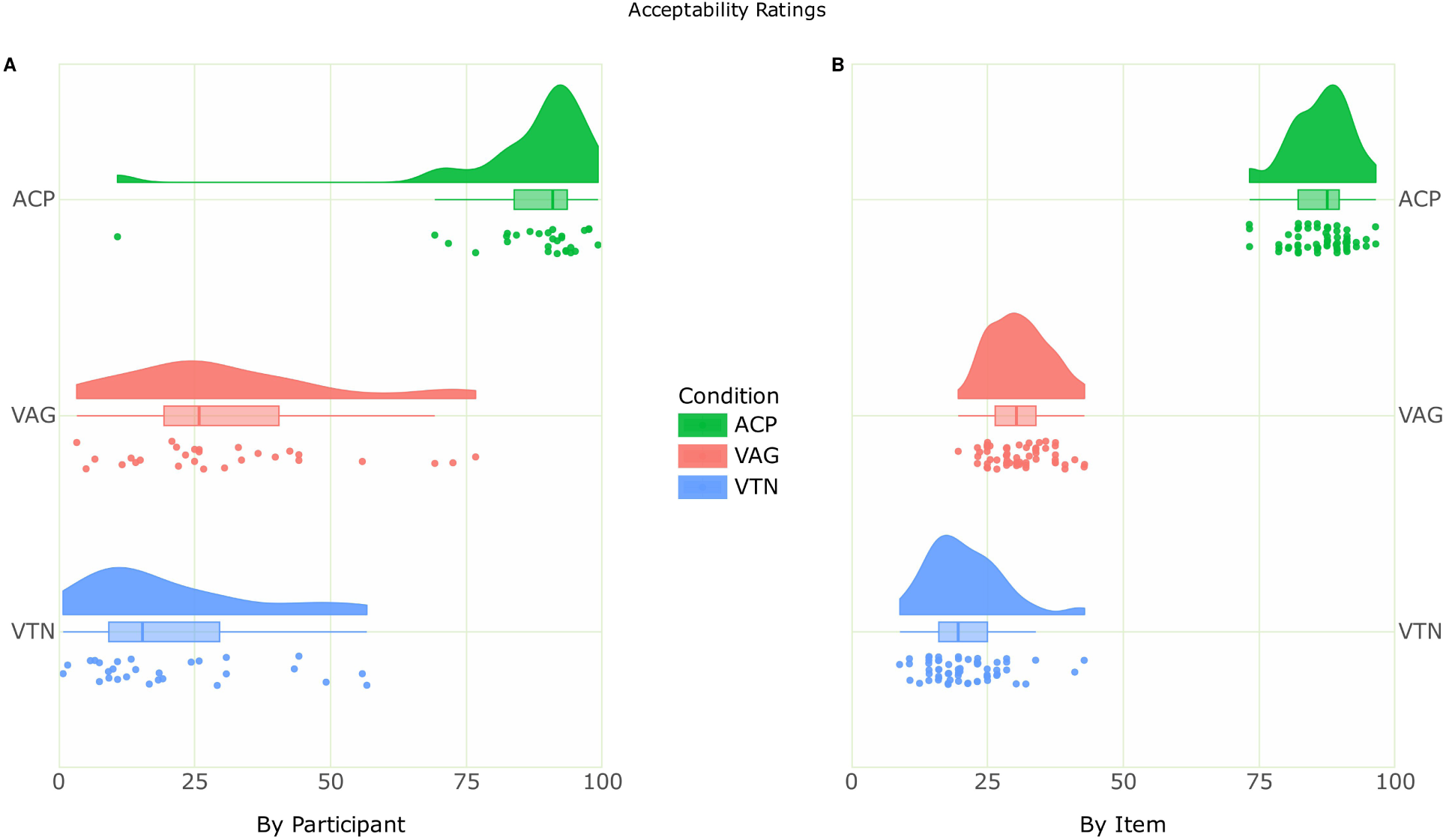
Raincloud plot of the acceptability ratings. Panel A shows the by-participant variability of acceptability ratings, with the individual data points representing the mean by-participant acceptability of each condition (ACP = Acceptable; VAG = Violation of Agreement; VTN = Violation of Tense). Panel B shows the by-item variability of acceptability ratings, with the individual data points representing the mean by-item acceptability of each condition.

The behavioral acceptability and accuracy were analyzed by fitting generalized linear mixed models using the lme4 package in R. The analysis code and full model outputs are available as R notebooks in the data analysis repository online. Categorical fixed factors used sum contrasts. In the analysis of acceptability data, the statistical model included the fixed factor Condition, with random intercepts for participants and items, and by-participant random slopes for the effect of Condition. Type II Wald chi-square tests of the fitted model (AIC = 7703.59) of the acceptability data showed a main effect of Condition (χ^2^(2) = 183.59, p < 0.001, s = 132.43). Estimated marginal means on the response scale were computed using the emmeans package on the model to compute pairwise contrasts of estimates of the three conditions with the mvt adjustment applied for multiple comparisons. This revealed a simple effect of Condition for the contrast of the acceptable condition versus the agreement violation condition (estimate = 0.666, SE = 0.042, p < 0.001, s = ∞), for the contrast of the acceptable condition versus the tense violation condition (estimate = 0.775, SE = 0.032, p < 0.001, s = ∞) , and for the contrast of the agreement violation condition versus the tense violation condition (estimate = 0.108, SE = 0.034, p = 0.004, s = 7.78). In the analysis of the probe detection accuracy, the statistical model included the fixed factor Condition, with random intercepts for participants and items. Models with complex random effects structures were singular. Type II Wald chi-square tests of the fitted model (AIC = 4904.34) of the accuracy data showed a main effect of Condition (χ^2^(2) = 14.23, p < 0.001, s = 10.27). Resolving this effect by computing estimated marginal means on the response scale with the mvt adjustment applied for multiple comparisons showed a simple effect of Condition for the pairwise contrast of estimates of the acceptable condition versus the tense violation condition (estimate = 0.022, SE = 0.006, p = 0.002, s = 8.73).

### 5.2 ERP Data

The ERPs at the verb are shown in Figure 3 for the critical conditions, with the difference plots showing the topography of effects for the two violation conditions compared to the acceptable condition in the two time-windows selected for analysis, namely 350-500 ms and 650-850 ms. These time-windows were chosen based on visual inspection and the typical latencies of ERP components that are known to be relevant for processing agreement.

**Figure 3:**
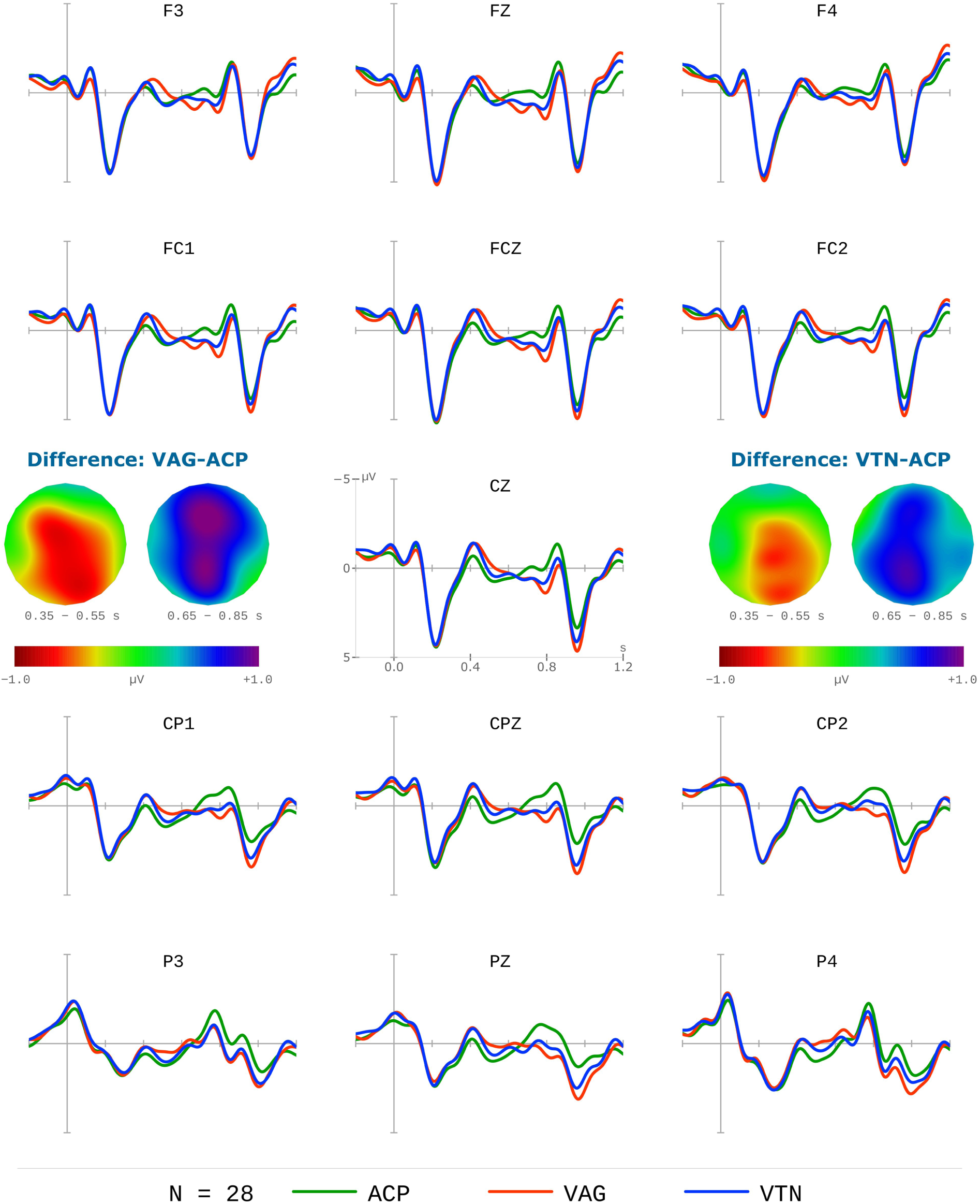
Grand-average ERPs at the verb for the three critical conditions (ACP = Acceptable; VAG = Violation of Agreement; VTN = Violation of Tense) from 28 participants. Negativity is plotted upwards, and the time axis runs from -0.2 s to 1.2 s (i.e., -200 ms to 1200 ms) with 0 being the onset of the critical verb. The difference plots show the topography of the negativity and late-positivity effects in the two time-windows (350-500 ms, 650-850 ms) for the agreement violation condition (left) and the tense violation condition (right) as opposed to the acceptable condition.

#### 5.2.1 Time-window 350−550 ms

The predominant effect in this time-window is the negativity for the agreement and tense violation conditions as opposed to the acceptable condition. Notably, there is virtually no difference between the agreement and tense violation conditions. The statistical linear-mixed model included the fixed factors Condition, ROI, the -200-0 ms pre-stimulus baseline mean amplitude as a covariate (scaled and centered), with random intercepts for participants, and by-participant random slopes for the effect of Condition. Type II Wald chi-square tests of the fitted model (AIC = 827.63) of the ERP data in the lateral regions showed a main effect of Condition (χ^2^(2) = 16.54, p < 0.001, s = 11.93). Estimated marginal means on the response scale were computed using the emmeans package on the model to compute pairwise contrasts of estimates of the three conditions with the mvt adjustment applied for multiple comparisons. This revealed a simple effect of Condition for the contrast of the acceptable condition versus the agreement violation condition (estimate = 0.400, SE = 0.102, p = 0.001, s = 9.71), and for the contrast of the acceptable condition versus the tense violation condition (estimate = 0.242, SE = 0.099, p = 0.05, s = 4.26). Type II Wald chi-square tests of the fitted model (AIC =1208.45) of the ERP data in the midline regions showed a main effect of Condition (χ^2^(2) = 27.95, p < 0.001, s = 20.15). Estimated marginal means on the response scale were computed using the emmeans package on the model to compute pairwise contrasts of estimates of the three conditions with the mvt adjustment applied for multiple comparisons. This revealed a simple effect of Condition for the contrast of the acceptable condition versus the agreement violation condition (estimate = 0.679, SE = 0.128, p < 0.001, s = 14.37), and for the contrast of the acceptable condition versus the tense violation condition (estimate = 0.491, SE = 0.140, p = 0.004, s = 7.94).

#### 5.2.2 Time-window 650−850 ms

The predominant effect in this time-window is the positivity for the agreement and tense violation conditions as opposed to the acceptable condition. Again, there is virtually no difference between the agreement and tense violation conditions. The statistical linear-mixed model included the fixed factors Condition, ROI, the -200-0 ms pre-stimulus baseline mean amplitude as a covariate (scaled and centered), with random intercepts for participants, and by-participant random slopes for the effect of Condition. Type II Wald chi-square tests of the fitted model (AIC = 815.95) of the ERP data in the lateral regions showed a main effect of Condition (χ^2^(2) = 13.40, p = 0.001, s = 9.66). Estimated marginal means on the response scale were computed using the emmeans package on the model to compute pairwise contrasts of estimates of the three conditions with the mvt adjustment applied for multiple comparisons. This revealed a simple effect of Condition for the contrast of the acceptable condition versus the agreement violation condition (estimate = -0.590, SE = 0.165, p = 0.003, s = 8.20), and for the contrast of the acceptable condition versus the tense violation condition (estimate = -0.468, SE = 0.169, p = 0.02, s = 5.31). Type II Wald chi-square tests of the fitted model (AIC =1230.93) of the ERP data in the midline regions showed a main effect of Condition (χ^2^(2) = 19.63, p < 0.001, s = 14.16). Estimated marginal means on the response scale were computed using the emmeans package on the model to compute pairwise contrasts of estimates of the three conditions with the mvt adjustment applied for multiple comparisons. This revealed a simple effect of Condition for the contrast of the acceptable condition versus the agreement violation condition (estimate = -0.981, SE = 0.229, p < 0.001, s = 10.85), and for the contrast of the acceptable condition versus the tense violation condition (estimate = -0.792, SE = 0.230, p = 0.004, s = 7.71).

## 6 Discussion

The present study used a violation paradigm to explore the neurophysiological correlates of the processing of tense and agreement in Modern Standard Arabic. Critical stimuli were either fully acceptable or constituted an agreement violation or a tense violation. Behavioral results showed that the anomalous conditions were consistently rated as unacceptable compared to the acceptable condition. ERP results at the verb showed that both agreement violations and tense violations elicited a biphasic pattern consisting of a centro-parietal negativity effect followed by a late positivity effect. Based on the latency and topography of these effects as well as the experimental conditions, the negativity effects in our study can be plausibly interpreted as instances of an N400 effect and the late-positivities as instances of a P600 effect. Of central importance in our findings is that the ERP responses elicited by tense and agreement violations were qualitatively similar. We take this as an indication that the neural mechanisms underlying the processing of these two grammatical categories in Arabic overlap to a very large extent, if not identical.

The N400 effect we observed for agreement violations is in line with several previous cross-linguistic findings, including previous studies on Arabic (Muralikrishnan & Idrissi, 2021; Idrissi et al., 2021; Bhattamishra et al., 2021; Zawiszewski, Santesteban, & Laka, 2016; Díaz, Sebastián-Gallés, Erdocia, Mueller, & Laka, 2011). The N400 effect elicited by tense violations is consistent with findings from previous studies on English (Allen, Badecker & Osterhout, 2003; Kutas & Hillyard, 1983). Whilst agreement violations have been traditionally viewed as formal morphosyntactic rule violations that engender LAN effects indicative of the detection of the morphosyntactic violation (Bornkessel & Schlesewsky, 2006; Friederici, 2002; Münte, Matzke, & Johannes, 1997; Münte, Szentkuti, Wieringa, Matzke, & Johannes, 1997), the likelihood of observing a LAN has been thought to be in direct proportion to the morphological richness of a language (Friederici & Weissenborn, 2007), and is said to depend upon whether morphological marking is crucial for assigning syntactic roles in the given language (Friederici, 2011). However, Molinaro, Barber, and Carreiras (2011) note that the nature of the complexities involved in morphological decomposition for feature identification may dissociate whether a LAN or, alternatively, an N400 ensues. Indeed, Choudhary et al. (2009) provide converging evidence from Hindi that this dissociation results from whether an interpretively relevant cue is violated (in which case an N400 ensues), or alternatively the violation involves a cue that is irrelevant for interpretation. In view of this, the absence of a LAN effect and the presence of an N400 in our study is not surprising, given that agreement computation in Arabic is not always predictable based on the features of the agreement controller alone, but is dependent upon specific syntactic properties of the construction involved, such as word-order and whether or not the subject is overt, as well as properties at the syntax-semantic interface such as humanness.

As for the P600 effect we observed for both types of violations, similar late-positivities have been reported in previous studies on tense and agreement processing. The P600 in our study can be interpreted as reflecting processes of repair or reanalysis associated with agreement violations (Bornkessel & Schlesewsky, 2006; Friederici, 2002, 2011) as well as structural anomalies (Osterhout & Mobley, 1995), and the effect is thought to be triggered by domain-general conflict monitoring processes (van de Meerendonk, Chwilla, & Kolk, 2013; van de Meerendonk, Kolk, Vissers & Chwilla, 2010; van de Meerendonk et al., 2009). In sum, our results suggest that violation of both tense and agreement in Arabic incur qualitatively similar repair or reanalysis costs, regardless of whether the violation involves temporal reference or agreement features.

Our initial rationale for conducting this study was to shed light on the nature of tense and agreement and the relationship between them in syntactic theory as well as the commonly reported dissociation between them in agrammatism. In the early generative syntactic tradition, Tense and Agreement were treated as distinct functional projections (see Pollock’s (1989) Split-Inflection Hypothesis). By contrast, in many later minimalist approaches, it was proposed that agreement features are encoded within the tense node itself (Chomsky, 2000). If these two categories were fundamentally different and represented in fundamentally different ways, one might expect to see qualitatively different neurophysiological signatures during their processing. However, our results point towards the exact opposite: the ERP patterns elicited by tense and agreement violations are identical, which suggests that the parser does not treat these features as being different during online sentence comprehension. We take our findings to be compatible with feature-based approaches in which tense and agreement form a single bundle of features rather than distinct structural projections or functional nodes (see the processing-based explanations of tense-agreement dissociations in agrammatism).

The results from our study also speak to the behavioral neuropsychological literature on agrammatic aphasia, in which the dissociations between tense and agreement have been central. Within the distinct projections approach, the Tree-Pruning Hypothesis (Friedmann & Grodzinsky, 1997; Friedmann, 2001) was proposed to account for this dissociation.

Specifically, the TPH relies on a syntactic structure showing different projections for tense and agreement where the former is hosted by a node higher than the latter. The observed dissociation between the two functional categories has been interpreted as reflecting damage to functional projections higher than Agr, which projections include Tense and Comp (see Figure 1). However, as we have just seen, the ERP results obtained in the current study from neurotypical Arabic speakers do not show any qualitative processing differences between tense and agreement whatsoever. We are therefore compelled to explain the dissociations reported in agrammatism, if any, by invoking factors other than hierarchical organization.

These dissociations can be attributed to task demands, production versus comprehension differences, or differential processing costs associated with specific features. In this respect, it is important to reiterate that many post-TPH studies of the tense and agreement in agrammatism reported inconsistent findings (see discussion in the introduction), suggesting that the reported dissociations might not be as robust as they were initially thought to be, and that agrammatic tense and agreement data is better explained by other factors than the position in a hierarchical syntactic structure. This view is consistent with many studies questioning strictly hierarchical explanations of tense–agreement dissociations and instead attributing the reported patterns to distinct feature processing mechanisms (Burchert et al., 2005).

Our study constitutes an important cross-linguistic contribution to the ERP literature on morphosyntactic processing. We provide evidence from Arabic, an understudied language, characterized by a rich inflectional morphological system and a typologically distinct agreement system (see Idrissi et al., 2021; Muralikrishnan and Idrissi 2021). In Arabic, tense and agreement are morphologically encoded in different ways, with tense being largely expressed by templatic variations and agreement by affixation. Despite the difference in morphological exponence between the two categories, the brain responses to their violations are virtually indistinguishable in our results. We wish to argue that the processing system may operate at a level of abstraction that does not heed the surface differences in the specific morphological exponence of grammatical features. We may add that the ERP effects observed in our data appear to be insensitive to the specific (formal) structural differences between tense and agreement and may, instead, reflect much more general neurophysiological / neurocognitive mechanisms deployed in coping with morphosyntactic integration, repair, and reanalysis during sentence comprehension.

More generally, in view of the fact that the current study is part of a broader research program investigating the neurocognitive processing of Arabic morphosyntax, it is crucial to note that the the biphasic pattern of results from the present study (N400-P600) are consistent with previous findings on agreement processing in Arabic (Idrissi et al., 2021; Muralikrishnan and Idrissi 2021). Features such as humanness play a central role in processing agreement in Arabic and interact in complex ways with the number feature due to the presence of deflected agreement (Idrissi et al., 2021). Further, whilst the relative salience of agreement features (person, number, gender) differentially modulates agreement processing for singular and plural subjects (Muralikrishnan and Idrissi, 2021), agreement and tense are processed qualitatively similarly (the present study), and no differences were observed between past-to-future tense violations and future-to-past tense violations (refer plot included in the data analysis repository online). Taken together, these studies suggest that morphosyntactic processing in Arabic is largely shaped by language-specific morphosyntactic properties as well as sociolinguistic factors associated with diglossia. In this regard, a potential limitation of the present study stems from the fact critical stimuli were in Standard Arabic, a language that is the mother tongue of no one. Since Arabic speakers seem to juggle with two grammatical systems, looking into how tense and agreement are processed in Spoken Arabic should provide valuable insight and offer valuable contribution to the literature. Therefore, an important avenue for future research is to investigate the interaction between Standard Arabic and Spoken Arabic.

In conclusion, our findings show that while the theoretical and some behavioral neuropsychological studies treat tense and agreement as distinct categories, the ERP responses to tense and agreement violations in Standard Arabic reported in the present paper seem to deploy identical, or at least, largely overlapping neurocognitive mechanisms. This is an interesting finding given the distinct theoretical status of these two categories in various syntactic models and, more intriguingly, their different realizations in Arabic morphology.

We take this to indicate that the processing system may treat tense and agreement as components within a unified system, probably as features within the same feature bundle, rather than as distinct morphosyntactic operations manipulating distinct projections in a hierarchical syntactic representation.

More broadly, we maintain that the results reported in this study highlight the value of different types of so-called external evidence in informing theoretical debates about linguistic representations and the overall architecture of linguistic knowledge (see Prunet et al., 2000; Idrissi & Kehayia, 2004; Idrissi et al., 2008; Idrissi, 2018; Idrissi et al., 2021; Idrissi et al., 2025, for such studies in Arabic). In addition to child language data and data from brain damaged populations, ERPs collected from neurotypical adult brains offer an excellent window on how language processing is accomplished in real time and may be used to support or refute theoretical models.

## Data analysis repository

The statistical analysis code and full model outputs of all the analyses reported are available as R notebooks here: https://doi.org/10.5281/zenodo.19022850

## Acknowledgements

The research reported here conducted in the context of several research visits by R. Muralikrishnan was partly supported by a Research Institute grant (G1001) from New York University in Abu Dhabi to Prof. Alec Marantz, who we should like to thank here.

Ali Idrissi was at the United Arab Emirates University in Al Ain during the study.

